# LPT: Orchestrating bounded agentic loops for automated protein binder design

**DOI:** 10.64898/2026.10.29.762730

**Authors:** Michael Uckelmann

## Abstract

Current large language models (LLMs) can reason over scientific information and use code-based tools, but their training knowledge is broad rather than specialized, can be out of date and is hard to attribute to sources. Furthermore, unconstrained agentic workflows and unlimited tool use can consume large amounts of tokens. Here we present LPT (little protein tiger), a framework for fixed orchestration of LLM stages, each running a bounded agentic loop with custom system prompts and tool access, to enable de novo protein binder design from a single natural language prompt. The LLM stages can query a curated literature database, containing ∼14000 papers digested into key findings, enabling recall of specialized information. An LLM with database access answered 41/48 test questions drawn from database key findings correctly, while a model without access answered none. In three end-to-end design campaigns, initiated from a single prompt, LPT autonomously identified the design intent, selected targets, structures and hotspot residues and designed candidate binders to disrupt CALCRL/RAMP1, PD-1/PD-L1 and PVR-TIGIT interactions. Cross-validation with multiple folding models confirms a high in silico success rate at minimal token consumption and low computational costs.

## Introduction

Large language models (LLMs) have rapidly improved over the past years, to the extent that they are now increasingly used for knowledge-based scientific work involving information retrieval, analysis and reasoning over experimental data^1–6^. Providing models with access to deterministic computational tools is a particularly powerful approach to combine the models reasoning capabilities with reliable computations^7–11^. Some limitations remain. A model’s knowledge is fixed by data available at the training cutoff date. Training knowledge embedded in the parameters of general purpose models is necessarily very broad and accurate detailed facts (e.g. affinities between two proteins) cannot be recalled reliably from training knowledge alone. Models occasionally “hallucinate” and do not always assign the correct source to information. Especially in the scientific domain accurate reasoning over specialist knowledge is key, small details can matter substantially and assigning provenance is important for credibility. Providing LLM agents with access to databases holding domain-specific scientific knowledge can ground the models responses in established fact. Giving models access to code-based programmatical tools can help automate routine tasks with deterministic and reproducible results^2,5,7–10^. Recent attempts at building specialized LLM frameworks for the scientific domain show that combining deterministic code, accurate databases and reasoning capabilities of LLMs can significantly improve the model’s performance at scientific tasks^1–4,6,8^.

In parallel to the advances in LLMs, accuracy and speed of diffusion models for protein structure prediction and protein structure generation (protein design) improved substantially^12–21^. These models now enable generation of protein binders for a given target protein with a relatively high experimental success rate. Frameworks such as Bindcraft^19^, ProteinHunter^20^ or NISE^22^ consolidate multi-step design tasks using multiple design tools into reproducible and accessible pipelines. The recently released chaperone framework attempts to integrate multiple design tools using lightweight abstraction layers to improve interoperability and enable use by LLM agents^23^.

Here we explore if a structured framework, that provides LLMs access to a curated literature database and deterministic coding tools for protein structure analysis, enables reliable end-to-end automation of protein design tasks starting from natural language prompts.

## Results

The Little Protein Tiger (LPT) is a framework for automated protein design using fixed orchestration of LLM stages to design binders to disrupt disease-relevant protein-protein interactions. Each LLM stage has access to a set of tools written in python code (Table 1) and runs a bounded agentic loop within a narrowly defined task. The pipeline intentionally limits LLM freedom and urges use of code-based tools wherever possible with the aim of making the output more reliable and controlling token usage. The complete pipeline runs through different phases that take a user’s prompt, read the design intent, select an appropriate target protein, analyse the biology surrounding the target and select a protein complex structure to be disrupted. The structure is analysed and a protein design campaign is initiated to design mini-proteins or macrocyclic peptides meant to inhibit the targeted protein-protein interaction (Fig.1). The different LLM stages of the pipeline all have a distinct system prompt (”skill”, Supplemental Information 1) that defines their task and sets out stages to achieve it. Alongside the system prompt, either the user input or a structured handoff in JSON format from the previous stage is passed to the LLM.

**Table 1.**
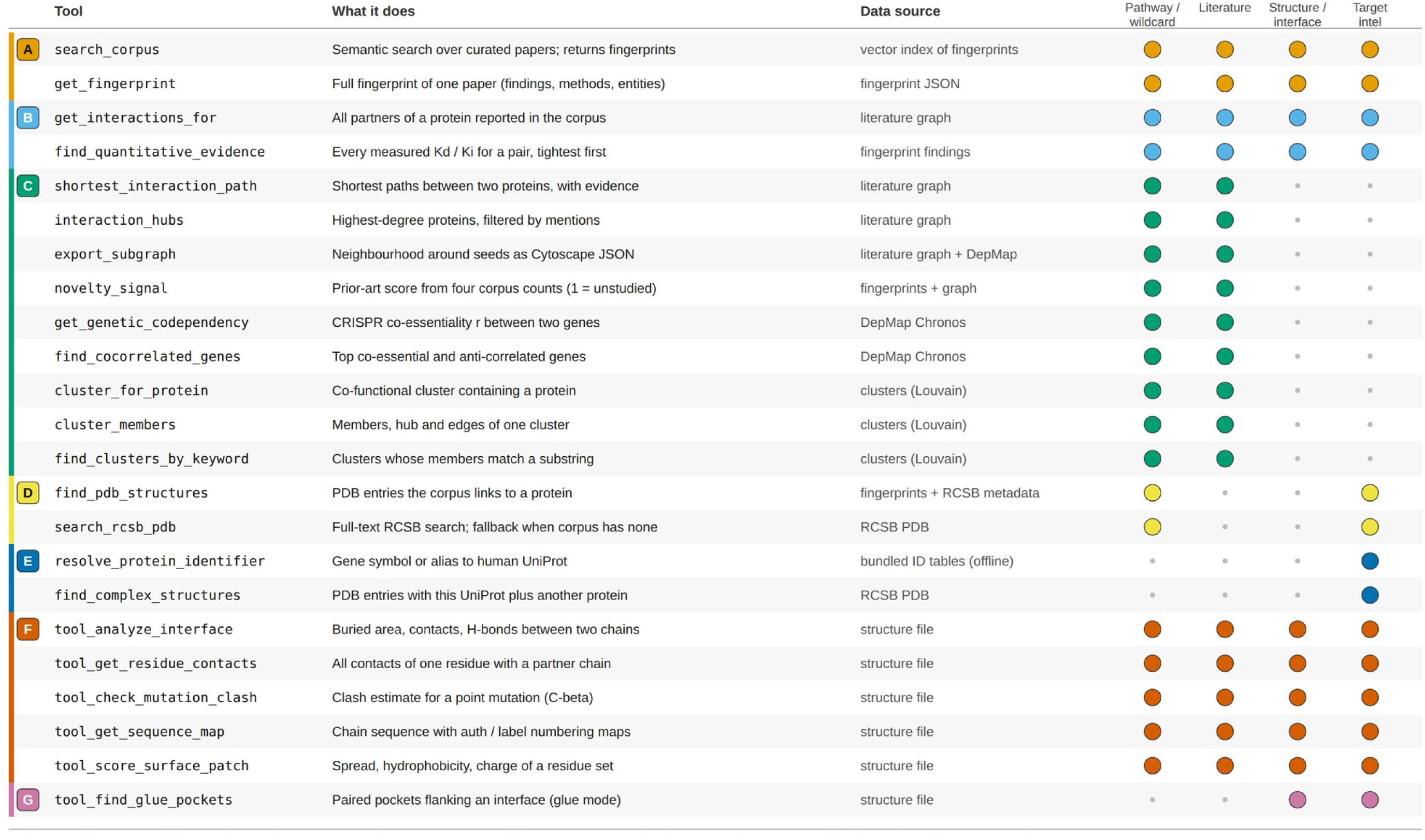
Tools available to the pipeline skills.

The target selection phase consists of two LLM stages, “pathway” and “literature”. Prompts to the target selection phase can be broad and should reflect the general design intent (e.g. “Design immune checkpoint inhibitors.”, Supplemental Information 2). The pathway-expert skill is tasked to understand the design intent and select an appropriate target. It has access to a custom-built literature database and tools to query the database. The second LLM stage uses the molecular-biology-expert skill as system prompt and is tasked with a more thorough analysis of the chosen target, the surrounding molecular pathways and currently available inhibitors. It suggests specific target sites and forwards a PDB structure of the target complex to the subsequent structure analysis phase. The complex-structure-analysis skill tasks the LLM to run a comprehensive protein structure analysis of a given PDB structure and determine the most suitable amino acid residues to be targeted (hotspot residues). The target decision along with the structure file and the chosen hotspot residues is then passed to the design phase. The design stage needs GPU access and runs diffusion models for protein design and protein structure prediction models for quality control and scoring. It starts with a small scale pilot, followed by a calibration run to size the production run scale. It then executes the production run at the determined scale, analyses results, ranks designs and provides a final summary of the campaign. In the following, the different modules of the pipeline and the literature database are discussed in detail.

**Figure 1:**
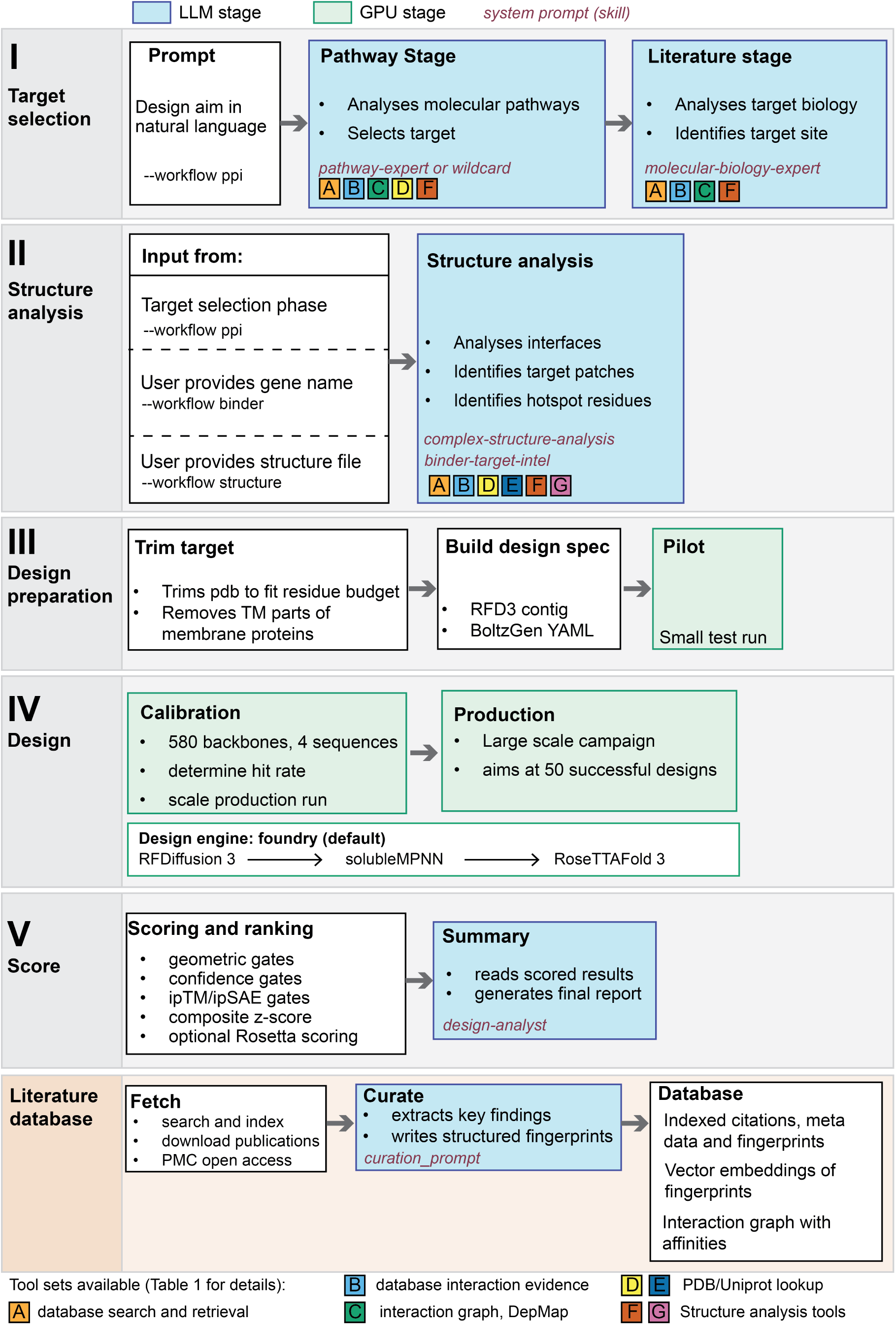
Flow chart illustrating the different stages of the pipeline. An end-to-end run is orchestrated by a pipeline runner and moves through all five stages automatically. Stages are resumable. The literature database is built independently and can be expanded.

### A literature database provides access to accurate scientific information

The pathway and literature stage have access to a literature database, built by curating scientific papers into fingerprints that hold key findings and quantitative information (Supplemental figure 3 and 4). The LLM stages are instructed to use this database as a source of information during reasoning. The database is constructed by user-defined keyword queries to NCBI PMC, Europe PMC and Semantic Scholar. Search results are indexed and stored in a SQLite database. For papers published in journals matching a user-defined journal tier list, the PDFs are downloaded, if available, with XML as a fallback option. To make paper contents accessible and searchable for the later LLM stages, each paper is curated into a fingerprint using an LLM curator. The curation prompt (Supplemental Information 3) is used as the system prompt in the LLM call, alongside the paper’s text. Only text and figure captions are considered, figures themselves are not processed. The LLM is tasked to understand the key findings from the paper and extract supportive quantitative and qualitative evidence for each key finding. It is instructed to assess negative and/or contradictory results, extract method details, PDB structure identifiers, determine which proteins interact according to the paper and find affinities. Finally, the curation skill writes a context hook summarizing the main findings of the paper (Supplemental information 4 for three example fingerprints^24–26^). The curation skill is tasked to emit the fingerprints in a structured JSON format with strict formatting guidelines. Guidelines are enforced via a Pydantic evaluation-correction loop during the curation call. The fingerprints are encoded into vectors using PubmedBERT embeddings^27^ and stored in a LanceDB vector database for semantic search. LLM skills can access the fingerprints via tools that query the vector database, the user can access the database directly via the conversational corpus-explorer tool. From all interacting protein pairs that have been identified across all papers in the literature database, an interaction graph is built. Proteins are the graph nodes. The edges carry mention counts (number of papers that mention the interaction) with supporting DOIs and all reported Kd and Ki values for the pair with citation and experimental context. The skills can further export a subgraph to which correlations of protein pairs across the DepMap CRISPR gene effect dataset^28^ are added to the edges (Fig.2A).

To assess if, and how, access to the literature database improves LLM responses we ran several ablation tests. We prompted the pathway expert, using Gemini Flash 3.7 as the model, with 7 different prompts in three different arms. The first arm used the standard pathway skill with access to all regular tools and the literature database. The second arm used a “wildcard” skill (Supplemental Information 1). This skill encourages the LLM to reason over information contained in the literature database and from DepMap correlations, as well as promoting more tool usage. In particular, the skill encourages to query the interaction graph and reason along edges (Fig.2A). The third arm resembles a straight LLM API call without any tool usage, the system prompt was an adapted pathway expert system prompt where all tool-use phases were replaced by one phase stating “no tools, work from your own knowledge” (full prompts and replies in Appendix I). Three replicates of the seven prompts were run for each setting to assess internal variability. Two different LLMs, claude-sonnet-5 and gpt-5.6-terra, were then tasked to judge the outputs according to a grading rubric. Each criterion was scored with a grade of 1-5 (1 = poor or wrong, 5 = excellent, Appendix I). We do not pose a verdict on the quality of the chosen targets and the judging models themselves might be steered by their training knowledge.

**Figure 2.**
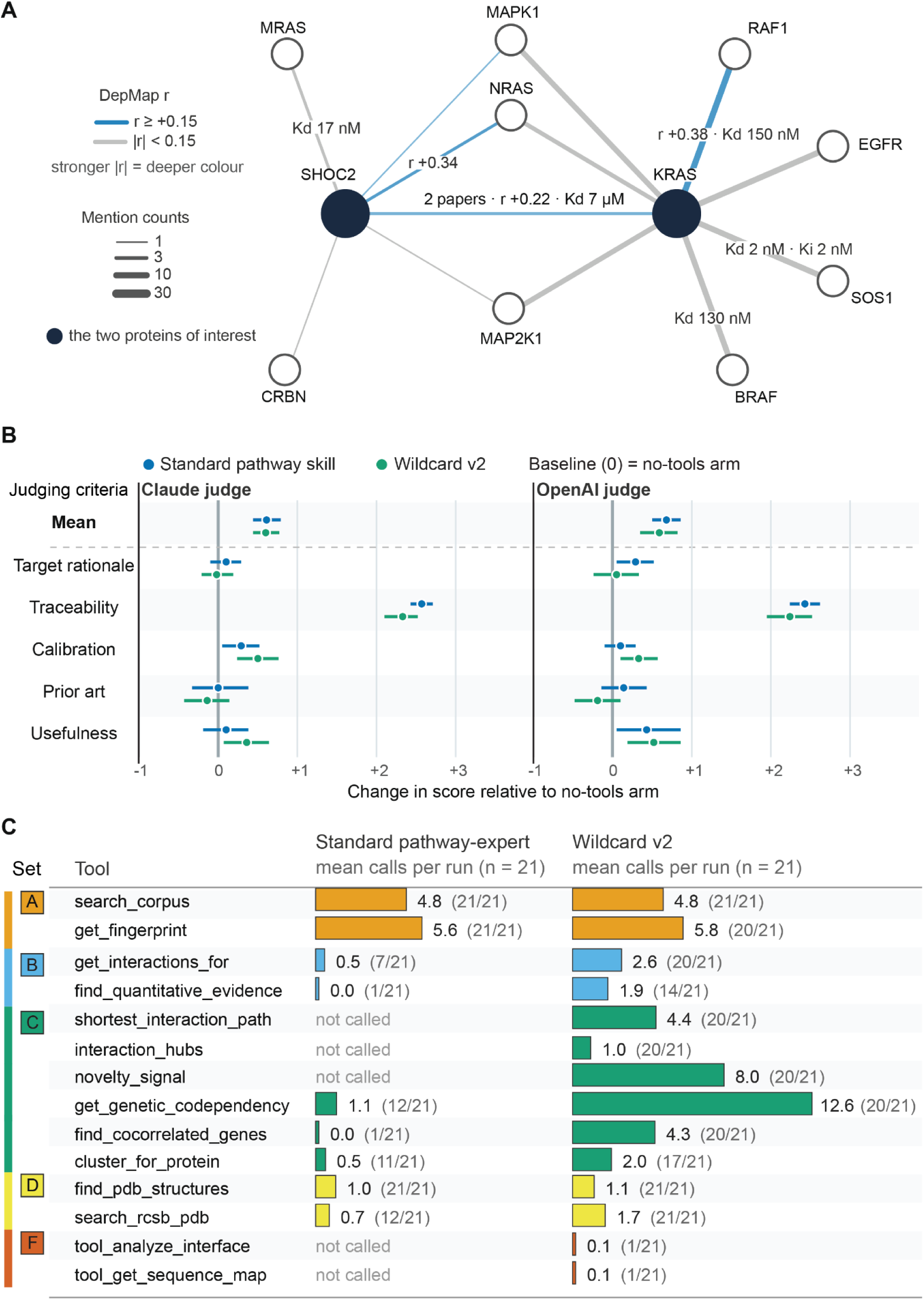
Ablation studies and different skills test the impact of the literature database. **A)** Rendering of the interaction graph that the LLMs can query via toolset C. Nodes are proteins, edges indicate interaction and carry affinity information and number of papers that mention this interaction in the database. The illustrated graph only shows a limited number of nodes. Edge thickness corresponds to the number of papers mentioning the interaction, colour hue corresponds to DepMap Pearson correlation of the pair across the DepMap CRISPR gene effect dataset^28^. The graph only considers protein pairs included in the database. **B)** Response scores for the standard pathway expert skill and the wildcard expert skill as judged by claude-sonnet-5 and gpt-5.6-terra. Scores are shown relative to a run where the LLM did not have access to any tools (no-tools arm). Seven different prompts were given, three replicates per prompt, for a total of 21 responses for each arm. Each point is averaged over a prompt’s repeats and then over all seven prompts. Bar indicates 95% bootstrap interval (3000 iterations, reports were resampled with replacement within each prompt and arm) **C)** Comparison of tool calls between the standard pathway expert skill and the wildcard skill. Mean calls per run and total runs where the respective tool was called are shown.

On target rational, calibration, prior art and usefulness the responses score similarly (Fig.2B), skills with access to tools score marginally better than the no-tools arm. Traceability is the one category where database access leads to substantial improvements. This is not unexpected, models in the arm without database access are asked to cite only if certain, but still shows that access to a database can improve provenance assignment.

There is substantial overlap in the targets chosen across the different trials (Table 2), which may suggest that suitable targets for these prompts can be picked straight from training knowledge or that training knowledge dominates decision making. The wildcard skill chose the most varied targets, reflecting its instructions. The extensive tool-calling of the wildcard skill (Fig.2C) indicates that the skill accesses information contained in the database frequently during reasoning. If this information improves target choice remains untested and we expect that a more comprehensive database would be necessary to take full advantage of the possibilities afforded by the tools.

**Table 2.**
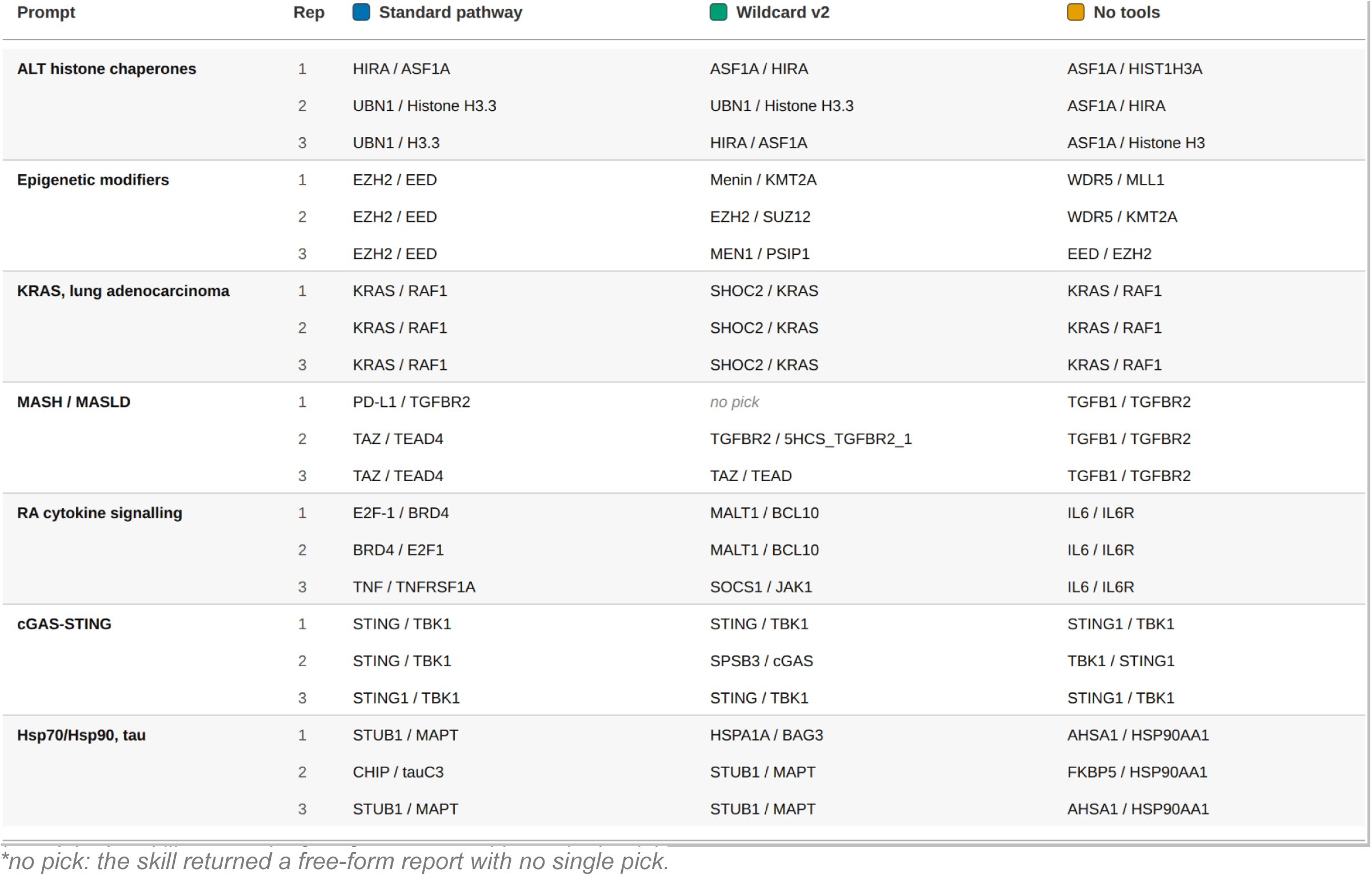
Target picks in the ablation studies.

To test if the LLM can retrieve detailed information from the literature database that is not immediately accessible through training knowledge, we ran closed-book and open-book exams using the corpus-explorer skill (Box 1), with Gemini-3.7-flash as the model. We chose 48 questions where the accurate answer is contained in the literature database fingerprints, including questions about specific affinities between proteins, specific residues involved in protein-protein interactions and about recent findings. In the open book case the expert skill had access to the LPT literature tools and database, in the closed book case not. The models were allowed to answer with “unknown” if they did not know the answer to a question. In the open book case the model answered correctly 41 times and answered “unknown” 7 times. In the closed book case the model never answered correctly, answered “unknown” 47 times and answered wrong once.

The complete exam with all questions and answers is provided in Appendix II. The test was necessarily biased towards information that was contained in the database and some questions were leading. Nonetheless, it shows that inclusion of a carefully curated literature database allows the LLM stages to find information not readily accessible from training knowledge.

In conclusion, the literature database serves as a knowledge backbone for the pipeline and models can recall detailed and accurate information from it. It has the potential to support reasoning based on quantitative evidence and across interaction networks.

#### Box 1

**Example questions and answers of the open/closed book test**

***“What dissociation constant (Kd) has been reported for the interaction between scFv6 and TcdB? Context: SPR analysis and cryo-EM structure determination of combinatorially assembled scFvs. Give one value with units.”***

*Answer: 72 nM. This value is reported from surface plasmon resonance (SPR) measurements of the *de novo* designed scFv6 binding to the Frizzled-binding epitope of *Clostridium difficile* toxin B (TcdB) in a study detailing antibody design using RFdiffusion (DOI: 10.1038/s41586-025-09721-5)*.

***A 2026 study examined ARX and PTPRD. Context: CRISPRi screen across 44 TFs. What did it report?***

*Answer: PTPRD is a convergent downstream effector gene regulated by ARX (and other transcription factors) that is enriched for neuropsychiatric disorder (schizophrenia/SCZ and major depressive disorder/MDD) risk genes (P < 0.05). This comes from a 2026 single-cell CRISPRi (Perturb-seq) screen evaluating 44 transcription factors in human radial glia during cortical neurogenesis (DOI: 10.1038/s41586-025-09997-7). Convergent downstream effectors like PTPRD showed significant enrichment for neuropsychiatric disease-*

### End-to-end orchestration of protein inhibitor design from a single prompt

The whole pipeline is best illustrated by following one specific example throughout all the stages. The examples shown here are taken from an actual run that proceeded end-to-end (Supplemental Information 5 for all generated stage reports). In this example case, the only input was one prompt:

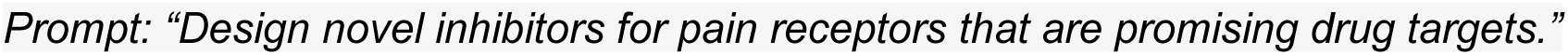

In the first stage of the pipeline, this prompt is passed via an API call to the chosen LLM (Gemini-3.7-flash in this case) along with the pathway-expert skill (Supplemental Information 1) as system prompt. The model is tasked to understand the disease context, pathways implicated in the disease, the disease mechanism and pathway dysregulation. It has access to all tools that interact with the literature database (Table 1). In the specific example, it categorizes the disease context as *“Chronic and inflammatory nociceptive pain”*, identifies key receptor-driven pathways involved and possible target nodes. The skill assessed the different target nodes for suitability and recommended to disrupt CALCRL-RAMP1 interaction. The decision was based on validated efficacy in pain and neurovascular inflammation, presence of a solvent-exposed extracellular domain and a high-quality experimental structure of the complex (Supplemental Information 5). Claims are supported by accurate citations from the literature database throughout^29–40^. A structured handoff with the target choice, structure suggestion and alternative targets is passed to the next phase. In between the first two stages the pipeline runs a deterministic step that is meant to improve structure choice. It collects alternative structures from the PDB, then compares them against the original suggestion of the pathway-expert skill. It triggers a switch if a better alternative is found. In the example a switch to a smaller, higher resolution crystal structure was triggered.

Stage two accepts the handover and runs the molecular-biology-expert skill. This skill is tasked with a more detailed analysis of the chosen target and the surrounding biology, specifically drawing from information contained in the literature database, with access to all literature tools. It is further asked to analyse known inhibitors and prior art, identify potential target sites from literature, including key residues, and provide a feasibility assessment. In our specific example, it analysed the RAMP1-CALCRL heterodimer role in CGRP-dependent pain signalling, highlighted known small molecule, antibody and small peptide inhibitors and identified key residues in the heterodimer interface that should be targeted. It accurately cites supporting evidence drawn from the literature database throughout^31,32,34,36–38^. The structured handoff for the next stage includes the PDB accession code of the selected structure and the suggested residues that should be targeted.

Stage three then runs an analysis of the protein structure to determine the final target site and hotspot residues that get passed to the design phase (Fig.3). A code-based tool is run before the LLM stage to prepare the structure. The tool downloads the structure file (if not on disk) and checks the chain identities. It generates a chain table that includes entity description and protein residue counts for each chain, computes buried surface area (BSA) and estimates H-bonds for every chain pair that involves the target protein. This information then gets combined with the handoff from the previous stage into a prompt for the subsequent LLM stage.

The combined prompt gets passed to the structure analysis LLM stage, with the complex-structure-analysis skill as the system prompt. The skill runs through an agentic loop, calling different tools to analyse the given complex structure. In a first phase the tool calls return an analysis of the interaction interface including residue- and interface-level BSA, H-bonds between interface residues and per-chain interface residues with per-residue information: residue type, hydrophobicity, contacts and an estimated binding contribution ***εε***. The model then groups residues into clusters based on shared contacts (Fig.3A and B).

Phase two calls a tool that analyses the clusters and returns hydrophobic fraction, spatial spread of the residues and a suitability rating. The model then chooses the most suitable target patch (tight clustering, hydrophobic anchors, disrupts target interface) and runs a literature cross-check against the literature database to assess what, if anything, is known about the chosen hotspot residues. Finally the skill generates residue hotspot lists in the appropriate format for RFD3^12^ and boltzgen^14^. In the example case, the skill determined two suitable target regions and chose region 1, a 10-residue patch with high aromatic density that spans the core heterodimerization surface on RAMP1 (Fig. 3A).

**Figure 3.**
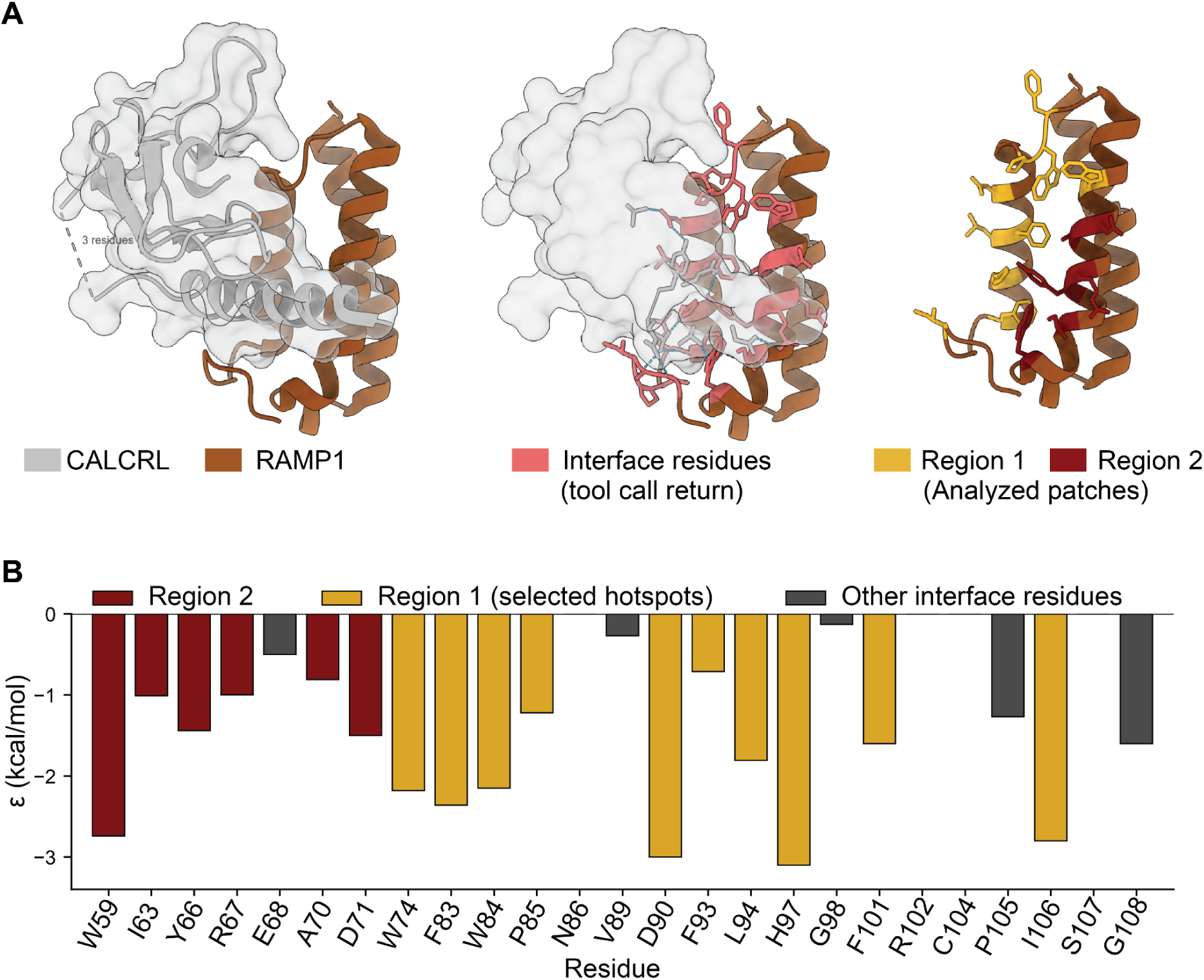
Target site selection during the structure analysis stage. **A)** The RAMP1-CALCRL complex structure selected for the design phase (left panel, PDB 3N7S^41^). The interface residues on RAMP1 identified by the structure analysis tool call are depicted in light red, H-bonding residues on CALCRL are shown as grey sticks (middle panel) and the target patches selected by the LLM phase are coloured in gold and maroon (right panel). **B)** The estimated binding contribution ***ε*** as returned by the structure analysis tool for each residue of the interface. Residues of Region 1 were forwarded as hotspots to the design phase.

The chosen hotspots are then passed to the actual design phase. Hotspot and chain information are translated into the appropriate input format for the chosen design backend, RFD3^12^ in the example case. All GPU-bound stages in the example case were run on a local workstation with a NVIDIA RTX Pro 4500 Blackwell GPU. LPT also supports staging of cluster runs and computing via Modal. Every design phase runs a 5-step program: (1) Diffusion with RFD3^12^; (2) A prefilter stage that removes designs with clashes, chain breaks and those with little secondary structure; (3) Sequence re-design with solubleMPNN^42^; (4) Refolding with RoseTTAFold3^43^; (5) Scoring of the refolds. This sequence is repeated at different scales: pilot, calibration and production. The pilot runs 100 designs to determine timing and prefilter pass rate. Calibration runs at a scale of 580 backbones, 4 sequences for each prefilter survivor, to determine hit rate for successful designs. Criteria for a successful design are: iPTM > 0.7, no clashes, dock RMSD ≤ 5 Å, fold RMSD ≤ 2 Å, binder pLDDT ≥ 0.75, interface PAE ≤ 15 Å, hotspot engagement ≥ 0.75. Based on the calibration, the production run scale is set to reach 50 successful designs. In the example case the hit rate was particularly high, which triggered an automatic raise of the iPTM target to >0.85.

The example production run produced 1396 refolded designs, 250 passed all gates and were considered successful. Successful designs are then scored with a combination of a composite z-score, iPTM and iPSAE^44^ (Supplemental Information 6). An optional Rosetta^45,46^ stage can add Rosetta interface energy (ΔΔG), Rosetta contact molecular surface and a penalty for buried unsatisfied polar residues. Finally, designs are ranked considering sequence diversity in addition to the composite score. Top ranked designs from the example campaign and from two additional end-to-end runs show good hotspot engagement, cover the native interaction interface and have promising in silico scores (Fig.4).

The second end-to-end campaign was started with the prompt “*Design immune checkpoint inhibitors.*” The pipeline designed binders against PD-L1 to disrupt interaction with PD-1 (Figure 4B, Supplemental information 2). The third campaign prompted to “*Design immune checkpoint inhibitors that are not targeting PD1 or PD-L1 but could synergize with PD1 or PD-L1 inhibitors.*” LPT chose to disrupt the TIGIT (T-cell immunoreceptor with Ig and ITIM domains)-PVR (Poliovirus receptor, CD155) interaction (Fig.4C). The in silico success rates for designs to pass all hard gates and iPTM ≥ 0.7 was 17.9% for the RAMP1 campaign and 9.6% for the PD-L1 campaign and 10% for the TIGIT campaign. Costs for the campaigns were $0.33, $0.42 and $0.39 in LLM API credits and ∼11 h, ∼34 and ∼20 GPU hours on the local workstation, respectively. To cross-validate the in silico success, the top 20 designs according to the LPT ranking were refolded in complex with the target using two independent folding models, Boltz-2^15^ and Protenix-v2^47^, from sequence only. The majority of the top 20 ranks in each campaign pass all gates in all folding models and show consistently high ipSAE scores (Table 3-5).

**Table 3.**
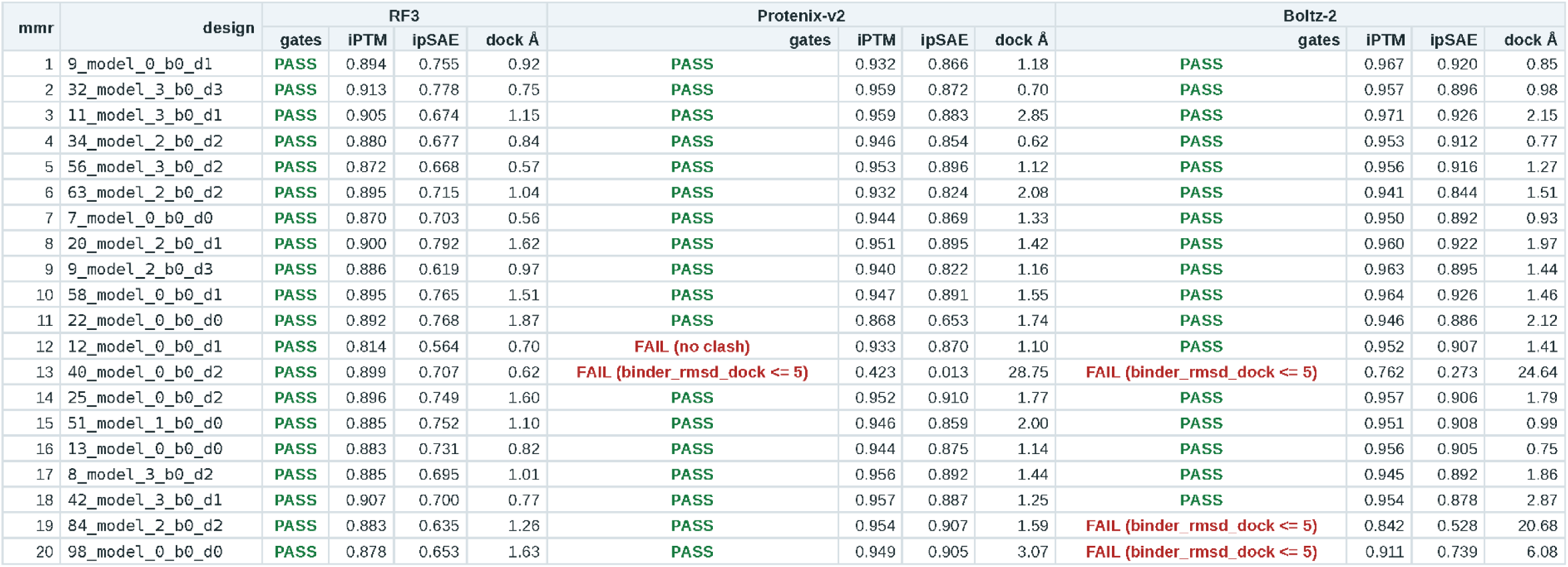
Cross-validation with Boltz-2 and Protenix-v2 of the RAMP1 campaign.

**Table 4.**
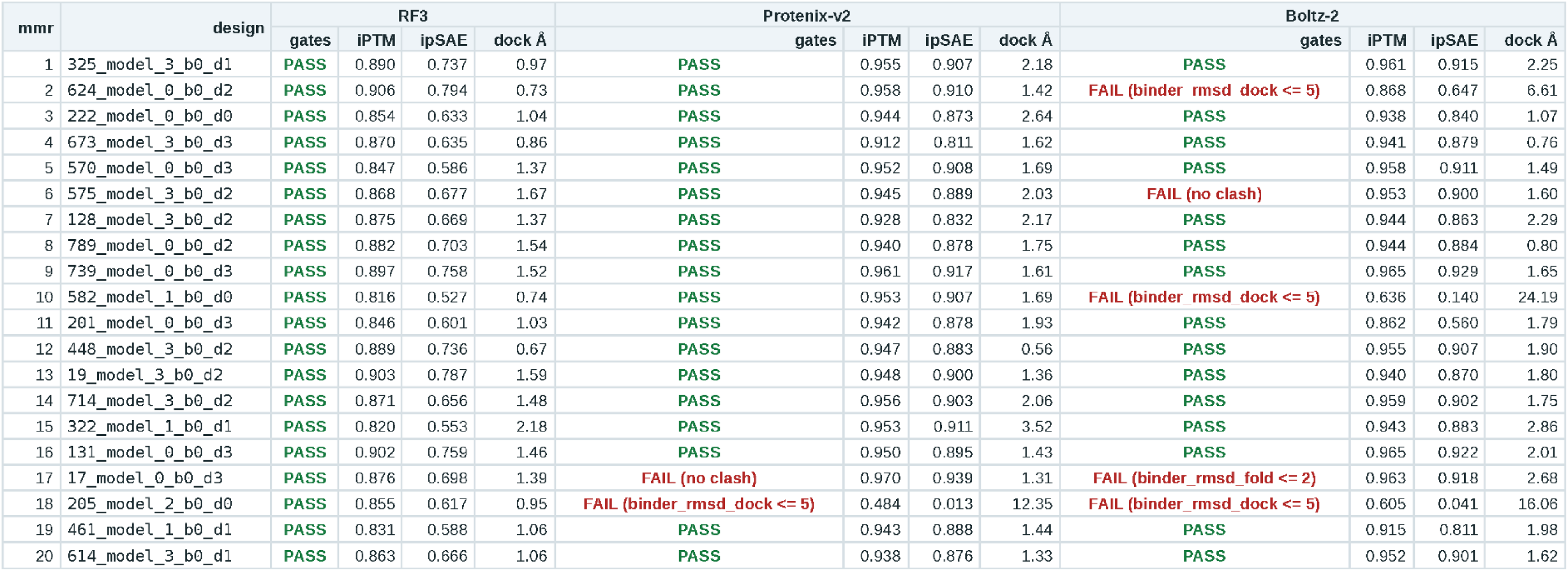
Cross-validation with Boltz-2 and Protenix-v2 of the PD-L1 campaign.

**Table 5.**
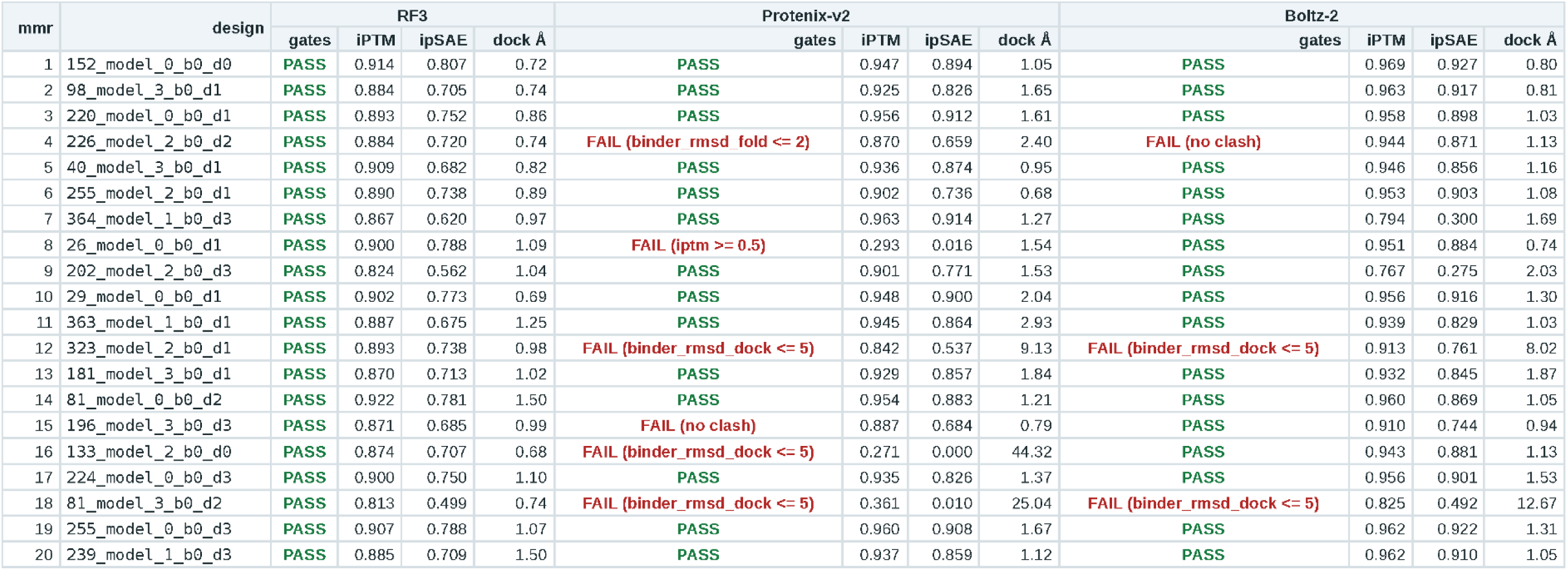
Cross-validation with Boltz-2 and Protenix-v2 of the TIGIT campaign.

### Alternative entry points

Next to end-to-end runs starting from one natural language prompt, the pipeline also supports alternative entry points. If the user knows which target to design against, they can provide the gene name as a query and enter the pipeline via the flag “--workflow binder”. The LLM stages for target selection and literature analysis are skipped in this case and the pipeline moves immediately to picking a structure and analysing it. The user can also provide a structure file directly, in which case the pipeline starts with analysing this structure. GPU stages remain the same across all entry points. The LLM skills can be run in isolation via the CLI with user-defined prompts. The literature database can be queried directly via a conversational LLM loop.

**Figure 4:**
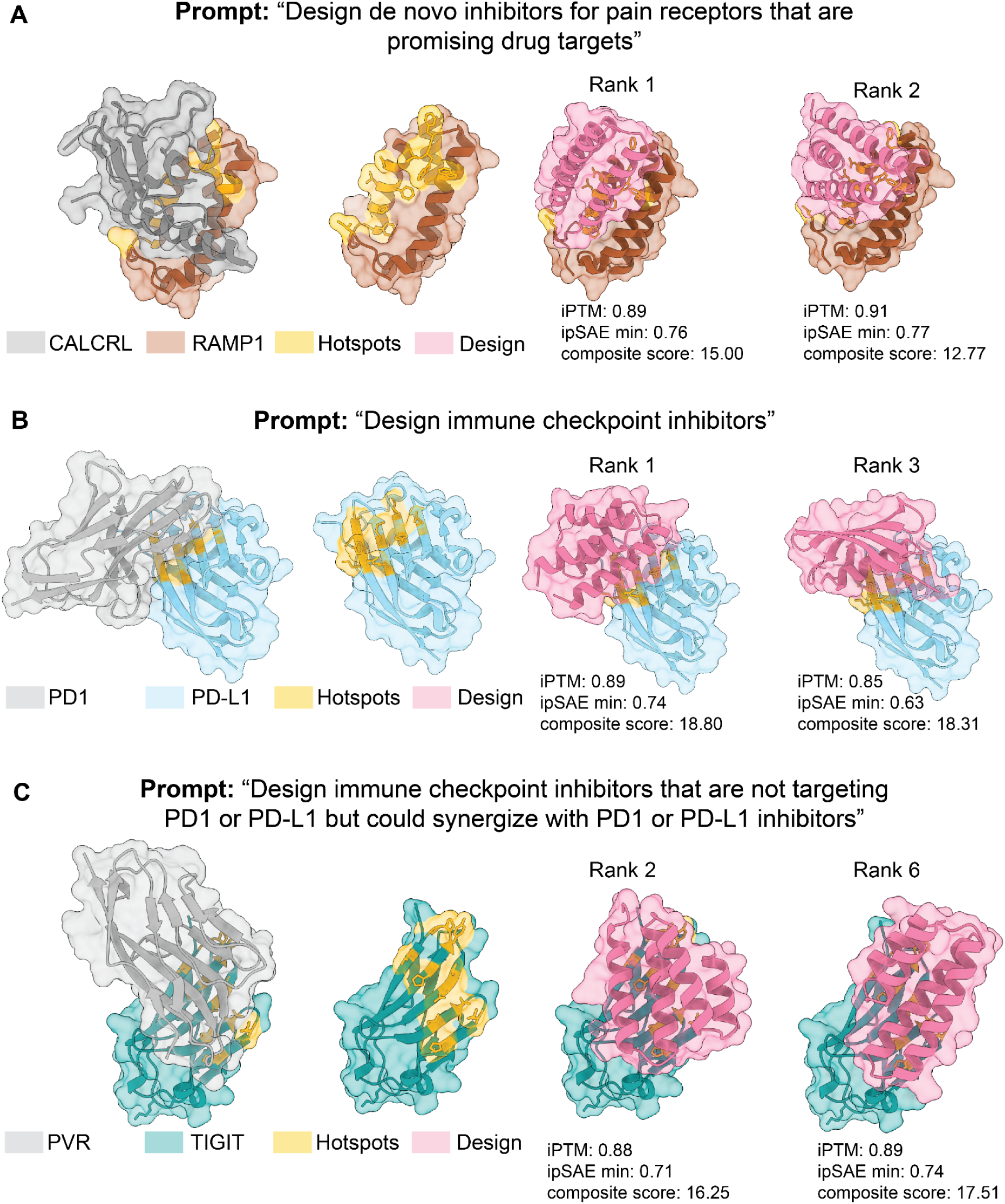
Binders generated from a single natural language prompt that pass in silico filters. **A)** Results of the pain receptor prompt end-to-end run. The two leftmost structures show the complex to be disrupted (PDB 3N7S^41^) and the model-chosen hotspots (gold). The two structures on the right show selected ranked designs. **B)** As in **A** but starting from the immune checkpoint prompt, the chosen target is a PD-L1/PD-1 complex (PDB 4ZQK^48^). **C)** as in **A** starting from the immune synergy prompt. The chosen target is a PVR-TIGIT complex (PDB 3UDW^49^)

## Discussion and limitations

The presented framework illustrates how integrating LLM reasoning with custom tools for protein design, backed by a literature database, can successfully drive an automated protein design campaign aiming to disrupt disease-relevant protein-protein interactions. Multiple recent efforts have proposed LLM-based workflows to automate and accelerate scientific discovery^1,2,4,6,8,23,50^ and the recently released protein design model RFD4-Proteina^51^ was built with agentic workflows in mind. Anthropic has shown that Claude, with unlimited token usage and a GPU budget of $10000, can design binders against multiple different disease-relevant targets^52^. Our own end-to-end campaigns using LPT cost $0.33, $0.42 and $0.39 in LLM API credits and ∼11 h, ∼34 and ∼20 GPU hours using a local workstation (pain receptor prompt, immune checkpoint campaign and immune synergy prompt, respectively). We have not evaluated the experimental success rate for our pipeline and can only provide in silico metrics as a proxy for success, the published success rates for RFDiffusion with similar in silico filters appears to be high^12,13^. Somewhat surprisingly, we noticed that safety refusals were triggered when we attempted to use our pipeline with Claude Sonnet to design against three of the targets that Anthropic used in their campaign^52^, suggesting the published approach was not readily reproducible with the publicly available models when we tested.

The literature knowledge backbone of our pipeline is limited by the database size and we would expect the model’s target picks to improve and become more varied with access to a larger database. The example campaigns were run with a database size of ∼14000 papers, the released database is smaller because it is limited to papers with licences that explicitly allow redistribution of derivatives. The cost for paper curation with the default model was ∼$0.01 per paper, making expansion cost-efficient. Though we can’t comprehensively assess if the presence of the database leads to better target choices, it does enable access to detailed quantitative information that is not readily accessible from training knowledge. It also enables accurate tracking of sources and assigning provenance to claims.

Our pipeline is not reliant on any one LLM provider or model. Other models than the ones used can be integrated with minimal effort, but will affect costs and possibly output quality. Similarly, the diffusion and refolding backends can be swapped with little effort, especially if standardized communication frameworks between design software and LLM agents, like the recently developed chaperone^23^, become the norm. Automated protein design frameworks like the one presented here, combined with lab automation, have the potential to significantly accelerate hypothesis generation, testing and lead generation in the early stages of drug discovery.

## Code and data availability

The code is available at https://github.com/MichaUckelmann/little-protein-tiger, alongside data from the example campaigns.

## Acknowledgements

I want to thank Anastassis Perrakis, members of the Perrakis group and Xiaohu Guo for input and useful discussions in the early stages of the project and Danny Sahtoe and members of the Sahtoe group for input and discussions during the later stages.

## Declaration of LLM use

Claude Code was used to write python code and code documentation and suggested improvements to the system prompts.

## Methods

### Building and expanding the literature database

The literature database is built by searching NCBI PMC, Europe PMC and Semantic Scholar with user-defined keywords. The search results get indexed and stored with metadata in a SQLite database. Papers get assigned a priority score (score = min(1.0, 0.55 * journal_tier + 0.3 * pub_type + 0.15 * recency + 0.08 * citation_boost)). Journal tier is defined by a journal tier list that can be edited by the user. For papers published in journals that are mentioned in the journal tier list, PDF download is attempted from the AWS PMC open access bucket, with XML download as a fallback option. Downloaded papers can then be curated via an LLM skill (Supplemental Information 3) using the curate_papers.py function. By default Google’s gemini-3.1-flash-lite is used as the curation model. The PDF text is extracted with PyMuPDF and processed according to the stages outlined in the curation prompt. Figures are not considered. The requested output is a structured JSON fingerprint with pre-defined fields that include the papers key findings with associated evidence, quantitative data, protein-protein interactions measured, PDB codes, a list of protein pairs that interact according to the evidence and negative results. The fingerprint includes a context hook that summarizes the main findings in a short paragraph. Fingerprints are validated by Pydantic during the LLM curation phase and sent back to the model with a Pydantic error if correction is necessary. At the end of a curation run the interaction graph is rebuilt with the new protein interaction information. Fingerprints (excluding numerical fields) are embedded using the PubMedBERT^27^ language model into 768-dimensional L2 normalized vectors and stored in a LanceDB. The LLM stages can semantically search the vector database.

### Execution of LLM stages

Each LLM stage runs a provider-agnostic agentic loop (Google, Anthropic or OpenAI API). The stage’s skill definition (Supplemental Information 1) gets passed as the system prompt, the stage input (either handoff from a previous stage or direct user input) gets passed as the user prompt. Alongside system and user prompt, the model gets access to stage-specific tools to search and retrieve information from the literature database, for graph queries and for protein structure analysis (Table 1). If the model decides to call a tool, this tool gets executed locally and the returns are appended to the conversation. Multiple tool calls are possible (maximum 30 on default). When the model responds without a tool call, this is considered the final response and the stage ends. The final report ends in a structured handoff block. The full report gets saved to disk as a markdown file, the handoff block is parsed for deterministic checks and provided as input for downstream stages. If safety classifiers are triggered, the run ends with a note.

### Tools for LLMs to access the literature database

The LLM stages have access to the information contained in the literature database through various tools, which are described in brief in the following:

**Search_corpus.** Runs a semantic search over the fingerprint embeddings that are stored in the vector database. It returns the top-k papers by cosine similarity with their DOI, study category and key findings.

**Get_fingerprint.** Retrieves the whole fingerprint for a given paper.

**Get_interaction_for.** For a given protein, returns all interacting proteins reported in the literature database, alongside mention counts, supporting DOI, Ki and Kd values

**Find_quantitative_evidence.** For a given protein pair, returns every measured affinity contained in the literature database

**Shortest_interaction_path.** Queries the interaction graph built from reported interactions between proteins, based on the literature database. It takes two proteins as input, determines the shortest path between them along the graph, then returns the nodes and edges along that path

**Get_genetic_codependency.** If DepMap data is available, it returns the Pearson correlation of essentiality between two genes.

**Find_cocorrelated_genes.** Given an input gene, finds all genes co-correlated in the DepMap CRISPR gene effect dataset^28^ with a minimum |r| of 0.2 with data in at least 100 cell lines. Returns the top 25 co-correlated genes. The model can override the default parameters.

**find_pdb_structure.** Finds PDB accession codes for target proteins in the literature database.

**Search_rcsb_pdb.** Allows directly searching the RCSB PBD if no PDB accession is found in the literature database

### Tools for LLMs to analyze protein structures

The LLM agent of the structure stage has access to deterministic tools for protein structure analysis implemented with BioPython and Gemmi. These tools, if called by the LLM, execute locally on the downloaded mmCIF or PDB file. The tools return numerical output to the agent. A brief description of the key tools follows.

**Tool_analyze_interface.** The main tool for interface analysis. Skills are asked to only call it once per complex structure. Given a structure of a protein complex, it returns the following metrics:

**Buried surface area (BSA):** BSA is computed by determining the solvent accessible surface area (SASA) for the complex and for the target chains in isolation. SASA is determined by the Shrake-Rupley algorithm^53^, rolling a 1.4 Å probe over the surface with BioPython default van der Waals radii per atom, 100 test points per atom are summed to residue level. BSA for individual residues is then calculated as SASA of that residue on isolated chains minus the SASA of that residue in the complex structure. Interface BSA is the sum of residue BSA of both chains across the interface.

**Interface residues and contacts:** Residues belong to an interface if any of the residue’s heavy atoms lie within 4.5 Å of a heavy atom on the other chain. Residue-residue contacts are identified using a k-d tree search (SciPy KDTree implementation with default parameters). For each interface residue pair the minimum atom-atom distance and an interaction type is reported. The interaction type is determined based on the residue identities. The possible interaction types are hydrogen bonds, electrostatic attractive or repulsive, hydrophobic, aromatic, polar and van der Waals.

**Hydrogen bonds:** H-bonds are estimated by distance between two candidate residues N, O or S heavy atoms and a rough geometry check. To be considered as forming an h-bond the heavy atoms need to be ≤ 3.2 Å apart and the angle D-A-X needs to be at least 90 degrees (D= donor, A = acceptor, X = the closest heavy atom covalently bonded to the acceptor).

This is an estimate of best candidates for hydrogen bonds rather than a rigorous test.

**Per-residue ΔΔG estimate *ε***: For each residue a heuristic ΔΔG, denoted as ***ε***, is estimated using the following terms: -0.028 kcal/mol per Å of buried surface area (only for hydrophobic and aromatic residues), -1.0 kcal/mol per H-bond and -0.5 kcal/mol per attractive electrostatic interaction. Residues below -2.0 kcal/mol are considered good hotspot candidates.

**pLDDT:** For input from structure predictions the pLDDT is read from the B-factor column and low-confidence parts are flagged.

**Tool_score_surface_patch:** Scores a given set of residues for its suitability as a target patch. The spatial spread of the residues is calculated based on C-alpha coordinates.

Hydrophobicity for the patch is reported as the mean Kyle-Doolittle value for the given patch. Each residue is assigned a class (aromatic, hydrophobic, charged, polar, other). The hydrophobic fraction is reported as the share of aromatic and hydrophobic residues over all residues of the patch. The suitability ranking is based on fixed cutoffs: “Excellent”: hydrophobic fraction ≥ 0.5, spread ≤ 10 Å and patch size at least 3 residues; “Good”: hydrophobic fraction ≥ 0.3, spread ≤ 15 Å; “marginal”: hydrophobic fraction ≥ 0.2. Everything else is returned as “poor”.

### End-to-end pipeline run

An end-to-end run of the whole pipeline is initiated by running a python script: run_pipeline.py --workflow ppi --query “<*user prompt>*” --project <PROJECT name>

After the CLI validates the input, the pipeline runner (pipeline_runner.py) starts up. The default settings are:

Provider: Gemini (gemini-3.8-flash currently, gemini-3.7-flash for most examples presented in the paper),

Pathway mode: standard (“wildcard” optional),

Design engine: foundry (RFD3, solubleMPNN, RF3)

Modality: mini-protein

Agentic loop limit: 30 tool turns

Per-call input ceiling: 150k tokens

The pipeline runner then executes a set of pre-defined phases in sequence (Fig.1), starting with target selection, followed by structure analysis and finishing with GPU-based design phases. The LLM stages all have a fixed token budget (150k tokens by default) and a fixed

API budget can be set as well (estimated from the API cost table in the config). By default, each LLM stage is limited to 30 turns of tool-calling. If a safety classifier is triggered, stages stop. The pipeline is resumable from completed stages, if LLM stages get interrupted within the stage, due to token limits, cost caps or turn limits, they need to be restarted from the beginning. If no structure to design against is found, the pipeline pauses, the user can then provide a structure and resume.

Communications between stages are handled via structured JSON handoffs, which serve as input to the next stage, alongside the system prompt of the respective stage.

### GPU-bound stages

The GPU stages for examples shown in this preprint were run on a local Nvidia RTX Pro 4500 Blackwell with 32 GB memory. The pipeline also supports Modal integration (Modal account necessary) and cluster staging.

The design stage defaults to RFD3^12^ -> solubleMPNN -> RF3^43^. Under default settings, RFD3 runs with diffusion_batch_size: 4, inference_sampler.step_scale=3, gamma_0=0.2, infer_ori_strategy: hotspots, is_non_loopy: true and dialect: 2; MPNN defaults are temperature 0.1, n_seq 4; RF3 runs with template: target, diffusion_batch_size: 1 and Seed 0. The GPU-bound design stage starts with a pilot at a default scale of 100 designs to test the input specifications. If successful, it proceeds to a calibration stage that generates 570 RFD3 designs, 4 sequences per design with solubleMPNN and refolds them with RF3. Calibration is used to time the run and measure hit rate. Hits are defined as designs that clear all hard gates:

- Dock RMSD at most 5.0 Å.
- Epitope recall at least 0.5.
- Hotspot engagement at least 0.75.
- Fold RMSD at most 2.0 Å.
- Binder pLDDT at least 0.75.
- iPTM at least 0.5.
- Interface PAE at most 15 Å.
- Minimum contact at least 1.8 Å (no clash).

Additionally a success metric needs to be achieved, by default set to iPTM > 0.7. The design aim is set to deliver 50 hits in the production stage and this aim must fit into the budget. The default budget is 5 campaign days (120 GPU hours).

Based on hit rate and timing the calibration stage returns a verdict:

SCALE_UP: At least 5 hits and pessimistic cost fits budget -> Production is sized to the full 50 design goal

SCALE_UP_PARTIAL: at least 5 hits, pessimistic cost exceeds budget by at maximum 1.5-fold -> campaign proceeds at reduced aim if at least 1/10 of the initial aim can be achieved within budget

ITERATE: Fewer than 5 hits or the budget buys less than 1/10 of the initial aim -> pipeline stops and prints options with estimated time and costs.

STOP: no hits

If the pipeline proceeds and the pessimistic timing estimate is under 48 hours, the campaign runs on the local GPU, otherwise a cluster run is staged (if staging is set up).

### Scoring

LPT scores designs according to several metrics, based on refolding model confidence, similarity of the refolded complex with the initial design and (optional) physical Rosetta metrics. ipSAE values are computed from the PAE matrix^44^, additional evaluated confidence metrics are ipTM, PTM and overall pLDDT. To evaluate binder dock RMSD, the design and refolded structures are superimposed on the target protein only and the binder C-alpha RMSD is measured in that frame. To evaluate binder fold consistency, the binder in the refold and design structures are superimposed and RMSD of the binder is calculated.

Epitope recall compares the target protein’s residues within 8 Å of the binder backbone in refold and design structure. Hotspot engagement measures the fraction of hotspot residues contacted in the refolds. Clashes are designated as heavy atom distances below 1.8 Å. Hard gates for a design to be considered successful are by default set at: dock RMSD ≤ 5 Å, fold RMSD ≤ 2 Å, binder pLDDT ≥ 0.75, iPTM ≥ 0.5, interface PAE ≤ 15 Å, epitope recall ≥ 0.5, hotspot engagement ≥ 0.75, and no clash.

For the final ranking a composite score is calculated (for details see Supplemental Information 6) for all designs passing the hard gates. Only the best scoring sequence for each backbone is advanced. To select diverse sequences a Maximal Marginal Relevance algorithm is run on the amino acid sequences of the designs. And a final ranking score is calculated from the composite score and the diversity score as: λ · normalised_composite − (1 − λ) · max_similarity_to_picked. λ is set to 0.5 by default. Designs with sequence similarity of >0.7 with higher ranking designs are dropped.

### Cross-validation with alternative folding models

For cross-validation we ran refolding of the top 20 ranked designs from each campaign with Boltz-2 and Protenix-v2. A pre-computed MSA was provided for the target protein, the designed binders were assigned a dummy MSA (no hits). One diffusion sample and a single seed were run for each model. iPTM, ipSAE and binder dock RMSD were calculated as in the regular pipeline. The cross-validation is not part of the LPT default pipeline.

