## Appendix II for "LPT: Orchestrating bounded agentic loops for automated protein binder design"

### Appendix II - Closed-book against open-book questions

### Closed-book against open-book questions

48 questions | 16 per type | 2 arms

Questions whose answers are stated in a fingerprint of a paper in the corpus. The same model (gemini-3.7-flash) answered each question twice: closed-book, with no tools, and open-book, with the corpus search tools. **The gold answer is what the corpus fingerprint states, not an independently verified fact.** Answers are verbatim; labels are re-derived from the stored answers with the scoring rules below.

#### RESULT

| QUESTION TYPE | CLOSED-BOOK CORRECT | OPEN-BOOK CORRECT |
|---|---|---|
| kd | 0 / 16 | 15 / 16 |
| residues | 0 / 16 | 14 / 16 |
| recent | 0 / 16 | 12 / 16 |

#### INSTRUCTIONS GIVEN TO THE MODEL, VERBATIM

Closed-book system prompt:

```
You answer specific questions about published biochemistry and molecular biology.
If you cannot determine the answer, reply with exactly UNKNOWN on the first line.
Otherwise put the answer on the first line (for a number give one value with its units; for residues give a
comma-separated list like R123, Y45), then at most two sentences saying where it comes from.
```

Open-book system prompt (adds one line):

```
You answer specific questions about published biochemistry and molecular biology.
If you cannot determine the answer, reply with exactly UNKNOWN on the first line.
Otherwise put the answer on the first line (for a number give one value with its units; for residues give a
comma-separated list like R123, Y45), then at most two sentences saying where it comes from.
You have tools that search a database of curated papers. Use them if they help.
```

#### SCORING

| | |
|---|---|
| `kd` | Correct if the answer's value is within 3-fold of the gold value (log10 difference at most log10 3). |
| `residues` | Correct if at least half of the gold residues appear in the answer's first line. |
| `recent` | Correct only if both blind judges (Claude, OpenAI) label it correct against the gold finding; any other combination counts as not correct. |

#### RECENT-FINDING JUDGE PROMPT, VERBATIM

```
You compare a model's answer with a gold finding from a paper. Labels: correct = the answer conveys the same
main finding; partly = overlaps but misses or changes something essential; incorrect = a different or
contradicting finding. Return only JSON: {"label": "correct|partly|incorrect", "reason": "<=30 words"}.
```

#### TYPE 1 · DISSOCIATION CONSTANTS

Correct if the answer's value is within 3-fold of the gold value (log10 difference at most log10 3).

**kd00 · What dissociation constant (Kd) has been reported for the interaction between 142F1 antibody and cLDLR? Context: BLI analysis of Fab binding to cLDLR. Give one value with units.**

gold 5.80e-09 M (doi 10.1021/jacs.5c15047, 2025)

**closed-book** · **unknown**
UNKNOWN

**open-book** · **unknown** · 5 tool calls
UNKNOWN

**kd01 · What dissociation constant (Kd) has been reported for the interaction between BCL7A-R11S and NCP? Context: EMSA assay comparing WT and mutant BCL7A. Give one value with units.**

gold 5.19e-07 M (doi 10.1093/nar/gkaf273, 2025)

**closed-book** · **unknown**
UNKNOWN

**open-book** · **correct** · 3 tool calls
519 nM

This value comes from an electrophoretic mobility shift assay (EMSA) measuring the binding of mutant BCL7A-R11S to the nucleosome core particle (NCP), as reported in the study on the structure of nucleosome-bound human BCL7A (DOI: 10.1093/nar/gkaf273).

**kd02 · What dissociation constant (Kd) has been reported for the interaction between Bebtelovimab and SARS-CoV-2 Spike? Context: Binding kinetic measurements across viral variants. Give one value with units.**

gold 7.90e-10 M (doi 10.1021/acs.jmedchem.3c01543, 2024)

**closed-book** · **wrong**
0.046 nM

This binding affinity (Kd) for Bebtelovimab (LY-CoV1404) binding to the SARS-CoV-2 wild-type (Wuhan-Hu-1) Spike RBD was determined via surface plasmon resonance (SPR) measurements reported by Westendorf et al. (Cell Reports, 2022). Across tested variants (including Alpha, Beta, Gamma, and Delta), reported Kd values consistently remained potent in the 0.046 to 0.075 nM range.

**open-book** · **correct** · 2 tool calls
790 pM

This value is reported in a review on antiviral protein-protein interaction inhibitors (DOI: 10.1021/acs.jmedchem.3c01543), which notes that Bebtelovimab (LY-CoV1404) binds SARS-CoV-2 variants of concern with Kd values ranging from 790 pM to 4 nM.

**kd03 · What dissociation constant (Kd) has been reported for the interaction between Bicyclic Peptide and TAGA hDNA? Context: Fluorescence quenching assay in PBS buffer. Give one value with units.**

gold 1.90e-06 M (doi 10.1021/acs.joc.5c01902, 2025)

**closed-book** · **unknown**
UNKNOWN

**open-book** · **correct** · 2 tool calls
1.9 µM

This value was determined via fluorescence quenching assays measuring the binding affinity of a bicyclic Arc repressor peptide analogue to the specific TAGA operator DNA box in PBS buffer (DOI: 10.1021/acs.joc.5c01902).

**kd04 · What dissociation constant (Kd) has been reported for the interaction between NS3aH1 and SDFTCSGWGTGWHMC? Context: BLI measurements of 'drop-out' peptides. Give one value with units.**

gold 1.90e-06 M (doi 10.1021/jacs.4c13561, 2025)

**closed-book** · **unknown**
UNKNOWN

**open-book** · **correct** · 2 tool calls
1.9 µM

This value was determined by bio-layer interferometry (BLI) for the unmodified "drop-out" peptide binding to NS3aH1 (DOI: 10.1021/jacs.4c13561). Bicyclization via the multipurpose linchpin resulted in a loss of binding affinity for this sequence.

**kd05 · What dissociation constant (Kd) has been reported for the interaction between Cucurbit[7]uril and Substrate? Context: ITC titration of CB7 with the aromatic aldehyde substrate. Give one value with units.**

gold 7.14e-07 M (doi 10.1039/d4sc04577g, 2024)

**closed-book** · **unknown**
UNKNOWN

**open-book** · **correct** · 2 tool calls
714 nM (or 7.14e-7 M)

This value is derived from an isothermal titration calorimetry (ITC) measurement yielding an association constant (Ka) of 1.40 × 10^6 M^-1 for the 1:1 binding between cucurbit[7]uril and the aromatic aldehyde substrate. It was reported by in vitro supramolecular catalysis studies investigating CB7-embedded gold nanoparticles (DOI: 10.1039/d4sc04577g).

**kd06 · What dissociation constant (Kd) has been reported for the interaction between 5HCS_TGFBR2_1 and TGFβRII? Context: Biolayer interferometry and luciferase reporter assay in HEK293 cells. Give one value with units.**

gold 1.00e-09 M (doi 10.1038/s41467-025-57192-z, 2025)

**closed-book** · **unknown**
UNKNOWN
I cannot determine the exact reported Kd value for this specific de novo construct.

**open-book** · **correct** · 3 tool calls
< 1 nM

This dissociation constant was measured by biolayer interferometry in a study designing de novo 5-helix concave scaffolds (5HCS) targeting immune receptors (DOI: 10.1038/s41467-025-57192-z). In the same study, 5HCS_TGFBR2_1 inhibited TGF-β signaling in HEK293 cells with an IC50 of 30.6 nM.

**kd07 · What dissociation constant (Kd) has been reported for the interaction between PfNPM and H2A-H2B? Context: ITC at 20 C in 300 mM NaCl buffer. Give one value with units.**

gold 1.00e-06 M (doi 10.1016/j.jbc.2025.108379, 2025)

**closed-book** · **unknown**
UNKNOWN

**open-book** · **correct** · 3 tool calls
1.00 µM

This dissociation constant was measured by isothermal titration calorimetry (ITC) at 20 °C in 300 mM NaCl buffer for the interaction between *Plasmodium falciparum* nucleoplasmin (PfNPM) and the H2A-H2B dimer (Kd = 1.00 µM for PfNPM A3, 1.006 µM for PfNPM A1).

**kd08 · What dissociation constant (Kd) has been reported for the interaction between scFv6 and TcdB? Context: SPR analysis and cryo-EM structure determination of combinatorially assembled scFvs. Give one value with units.**

gold 7.20e-08 M (doi 10.1038/s41586-025-09721-5, 2026)

**closed-book** · **unknown**
UNKNOWN

**open-book** · **correct** · 2 tool calls
72 nM

This value is reported from surface plasmon resonance (SPR) measurements of the *de novo* designed scFv6 binding to the Frizzled-binding epitope of *Clostridium difficile* toxin B (TcdB) in a study detailing antibody design using RFdiffusion (DOI: 10.1038/s41586-025-09721-5).

**kd09 · What dissociation constant (Kd) has been reported for the interaction between Cdc27 and PCNA? Context: In vitro ITC titration with recombinant SpPCNA. Give one value with units.**

gold 3.50e-06 M (doi 10.7554/elife.91461, 2024)

**closed-book** · **unknown**
UNKNOWN

**open-book** · **correct** · 2 tool calls
3.5 ± 0.3 µM

This dissociation constant was measured by isothermal titration calorimetry (ITC) using the PIP motif peptide from the *Schizosaccharomyces pombe* DNA polymerase delta subunit Cdc27 and recombinant SpPCNA (Mattiroli et al., *eLife*, 2024, DOI: 10.7554/eLife.91461).

**kd10 · What dissociation constant (Kd) has been reported for the interaction between VHH_flu_01 and Influenza haemagglutinin? Context: Cryo-EM structural determination of VHH bound to trimeric H1 haemagglutinin. Give one value with units.**

gold 7.80e-08 M (doi 10.1038/s41586-025-09721-5, 2026)

**closed-book** · **unknown**
UNKNOWN

**open-book** · **correct** · 7 tool calls
78 nM

This affinity is reported for the de novo-designed nanobody VHH_flu_01 targeting trimeric H1 haemagglutinin in a study demonstrating atomic-level accuracy of RFdiffusion-designed antibodies (DOI: 10.1038/s41586-025-09721-5).

**kd11 · What dissociation constant (Kd) has been reported for the interaction between MDM2 and Peptide? Context: BLI screening of 15 peptides against MDM2. Give one value with units.**

gold 6.50e-08 M (doi 10.1021/acschembio.5c00774, 2025)

**closed-book** · **unknown**
UNKNOWN

**open-book** · **correct** · 6 tool calls
65 nM

This value is reported as the strongest affinity in a 65–165 nM range among 7 out of 15 BindCraft-designed peptides that bound MDM2 in biolayer interferometry (BLI) screening assays. It originates from an evaluation of the generative design platform BindCraft for producing high-affinity peptide binders (DOI: 10.1021/acschembio.5c00774).

**kd12 · What dissociation constant (Kd) has been reported for the interaction between NbPTHR1 and PTHR1? Context: Competition binding assays and epitope footprinting. Give one value with units.**

gold 1.00e-09 M (doi 10.1038/s41467-024-49068-5, 2024)

**closed-book** · **unknown**
UNKNOWN

**open-book** · **correct** · 2 tool calls
1 nM
This dissociation constant is reported for the binding of NbPTHR1 to the extracellular domain of PTHR1 in competition binding and epitope footprinting experiments (DOI: 10.1038/s41467-024-49068-5).

**kd13 · What dissociation constant (Kd) has been reported for the interaction between OA and HOXB9? Context: SPR assay. Give one value with units.**

gold 7.69e-07 M (doi 10.1038/s41467-025-67083-y, 2025)

**closed-book** · **unknown**
UNKNOWN

**open-book** · **correct** · 4 tool calls
7.69e-7 M (769 nM)

This value comes from surface plasmon resonance (SPR) measurements showing direct binding between oleic acid (OA) and HOXB9, reported in a study on metabolic syndrome-driven endometrial cancer progression (DOI: 10.1038/s41467-025-67083-y).

**kd14 · What dissociation constant (Kd) has been reported for the interaction between EPHA2 and Compound 23? Context: SPR binding affinity measurement. Give one value with units.**

gold 1.20e-07 M (doi 10.1021/acs.jmedchem.4c00411, 2024)

**closed-book** · **unknown**
UNKNOWN

**open-book** · **correct** · 2 tool calls
120.4 nM

This value comes from surface plasmon resonance (SPR) binding affinity measurements of macrocyclic 2-aminopyrimidine kinase inhibitors binding to the EPHA2 kinase domain, reported by Wouters et al. (DOI: 10.1021/acs.jmedchem.4c00411).

**kd15 · What dissociation constant (Kd) has been reported for the interaction between LYN and CSK? Context: K562 cells treated with SI-3. Give one value with units.**

gold 3.46e-08 M (doi 10.1038/s41586-025-09763-9, 2026)

**closed-book** · **unknown**
UNKNOWN

**open-book** · **correct** · 3 tool calls
34.6 nM

This dissociation constant was measured in a study examining inhibitor-induced kinase destabilization in K562 cells treated with the CSK inhibitor SI-3 (DOI: 10.1038/s41586-025-09763-9).

#### TYPE 2 · INTERFACE RESIDUES

Correct if at least half of the gold residues appear in the answer's first line.

**rs16 · Which amino acid residues were reported as important for the interaction between PELP1 and SENP3? Context: In vitro proSUMO cleavage assay. List the residues.**

gold V776, E777, I778 (doi 10.1126/sciadv.adw4603, 2025)

**closed-book** · **unknown**
UNKNOWN

**open-book** · **correct** · 3 tool calls
V776, E777, I778
These residues are located within a short linear motif (residues 764–781) in the intrinsically disordered region of human PELP1 that mediates SENP3 binding and allosterically activates its endopeptidase activity in proSUMO cleavage assays (DOI: 10.1126/sciadv.adw4603).

**rs17 · Which amino acid residues were reported as important for the interaction between ECH and PAP? Context: X-ray crystallography of unliganded vs. liganded ECH. List the residues.**

gold Ala98, Gly141 (doi 10.1021/jacsau.6c00054, 2026)

**closed-book** · **unknown**
UNKNOWN

**open-book** · **correct** · 3 tool calls
A98, G141

These backbone amide nitrogen atoms of rat enoyl-CoA hydratase (ECH) form the oxyanion hole, which reorganizes and narrows from a 5.2 Å to a 4.0 Å N–N distance upon 3',5'-ADP (PAP) binding during crystallographic analysis (DOI: 10.1021/jacsau.6c00054).

**rs18 · Which amino acid residues were reported as important for the interaction between HalA and ODA? Context: Electrophoretic mobility shift assay (EMSA). List the residues.**

gold Q178, W27, E28, S141, N143 (doi 10.1038/s41586-025-09058-z, 2025)

**closed-book** · **unknown**
UNKNOWN

**open-book** · **correct** · 3 tool calls
W27, E28, S141, N143, Q178

These residues were identified in electrophoretic mobility shift assays (EMSAs) and structural analyses characterizing the binding of oligodeoxyadenylate (ODA) to the HalA crown domain in the bacterial Hailong anti-phage defense system (DOI: 10.1038/s41586-025-09058-z).

**rs19 · Which amino acid residues were reported as important for the interaction between YPEL5 and WDR26? Context: Structural comparison and in vitro ubiquitylation. List the residues.**

gold G2, R3, F5 (doi 10.1016/j.molcel.2024.04.014, 2024)

**closed-book** · **unknown**
UNKNOWN

**open-book** · **correct** · 2 tool calls
G2, R3, F5
These N-terminal residues of YPEL5 were identified as a basic degron mimic that binds WDR26 and competitively inhibits the ubiquitylation of substrates such as NMNAT1 by the CTLH E3 ligase complex. This mechanism was demonstrated through structural comparison by cryo-EM and in vitro ubiquitylation assays (DOI: 10.1016/j.molcel.2024.04.014).

**rs20 · Which amino acid residues were reported as important for the interaction between VPS34 and MSB? Context: Rescue assays in Pik3c3loxP/loxP MEFs. List the residues.**

gold Cys54, Cys61 (doi 10.1126/science.adk9167, 2024)

**closed-book** · **unknown**
UNKNOWN

**open-book** · **unknown** · 6 tool calls
UNKNOWN

**rs21 · Which amino acid residues were reported as important for the interaction between CDK2 and CDK1? Context: LANCE Ultra TR-FRET enzyme inhibition assay of generated lead compounds. List the residues.**

gold LEU83, GLU81, LYS33 (doi 10.1038/s41467-024-46569-1, 2024)

**closed-book** · **unknown**
UNKNOWN

**open-book** · **correct** · 4 tool calls
K33, E81, L83

These residues were identified as key interaction sites in a study using the pocket-based molecular diffusion model (PMDM) to generate CDK2 inhibitors with selectivity over CDK1, evaluated via LANCE Ultra TR-FRET assays (DOI: 10.1038/s41467-024-46569-1).

**rs22 · Which amino acid residues were reported as important for the interaction between BAF and Lamin A/C? Context: Reconstituted nucleosome with 24 bp linker DNA. List the residues.**

gold Lys486, His506 (doi 10.1038/s41467-025-56823-9, 2025)

**closed-book** · **unknown**
UNKNOWN
I cannot determine the specific residue list for this reconstituted nucleosome condition with certainty.

**open-book** · **correct** · 2 tool calls
K486, H506

These Lamin A/C residues within the immunoglobulin-like fold (IgF) domain were identified by cryo-EM as critical for direct DNA contacts and complex formation on nucleosomes reconstituted with 24 bp linker DNA. Disease-associated mutations at these positions (K486N and H506D) significantly reduce nucleosome binding in pull-down assays (DOI: 10.1038/s41467-025-56823-9).

**rs23 · Which amino acid residues were reported as important for the interaction between LacZ and Glycosyrin? Context: Cryo-EM structural resolution of LacZ-glycosyrin complex. List the residues.**

gold E538, W569 (doi 10.1126/science.adp2433, 2025)

**closed-book** · **unknown**
UNKNOWN
I cannot find any published cryo-EM study or biochemical report detailing the interaction between LacZ and a compound named glycosyrin.

**open-book** · **correct** · 2 tool calls
E538, W569

A 1.9 Å cryo-EM structure of the LacZ–glycosyrin complex revealed that glycosyrin binds the LacZ active site by forming key non-covalent interactions with catalytic glutamate E538 and conserved active-site tryptophan W569 (DOI: 10.1126/science.adp2433).

**rs24 · Which amino acid residues were reported as important for the interaction between ALK-3 and ALK-3bp? Context: SPR measurement of binding kinetics. List the residues.**

gold Gly65, Leu96, Thr94 (doi 10.1038/s41467-025-67866-3, 2026)

**closed-book** · **unknown**
UNKNOWN

**open-book** · **unknown** · 6 tool calls
UNKNOWN

**rs25 · Which amino acid residues were reported as important for the interaction between CUL5 and Tau? Context: Reconstitution in HEK cells and in vitro using recombinant proteins. List the residues.**

gold K83, K85, K106, K117, K122 (doi 10.1016/j.cell.2025.12.038, 2026)

**closed-book** · **unknown**
UNKNOWN

**open-book** · **correct** · 2 tool calls
K83, K85, K106, K117, K122

These lysine residues were identified as key sites for CRL5(SOCS4)-mediated ubiquitination and interaction with Tau in reconstitution assays using HEK cells and recombinant proteins (DOI: 10.1016/j.cell.2025.12.038).

**rs26 · Which amino acid residues were reported as important for the interaction between AD-tau and D-TLKIVWI? Context: Dot blot and TEM analysis of AD-tau disaggregation. List the residues.**

gold Leu2, Ile4, Trp6 (doi 10.1038/s41586-025-09244-z, 2025)

**closed-book** · **unknown**
UNKNOWN

**open-book** · **correct** · 2 tool calls
L2, I4, W6

These residues within the D-TLKIVWI peptide were reported as essential for interacting with and disassembling AD-tau fibrils based on dot blot and TEM disaggregation assays. The findings are reported by a study investigating the strain-relief disassembly mechanism of Alzheimer's disease tau fibrils by D-peptides (DOI: 10.1038/s41586-025-09244-z).

**rs27 · Which amino acid residues were reported as important for the interaction between HalB and ODA? Context: Crystal structure of HalB D21A/D23A mutant. List the residues.**

gold Y227, D21, D23, R13, R128 (doi 10.1038/s41586-025-09058-z, 2025)

**closed-book** · **unknown**
UNKNOWN

**open-book** · **correct** · 3 tool calls
Y227, D21, D23, R13, R128

These residues were identified in the crystal structure of the HalB D21A/D23A mutant characterizing the Hailong anti-phage defense system (DOI: 10.1038/s41586-025-09058-z). They mediate the self-primed synthesis and binding of oligodeoxyadenylate (ODA), with Y227 acting as the initiating priming residue.

**rs28 · Which amino acid residues were reported as important for the interaction between Pigeonpox PDE and cGAMP? Context: In vitro biochemical assay using recombinant protein. List the residues.**

gold H72, H167 (doi 10.1038/s41586-024-07809-y, 2024)

**closed-book** · **unknown**
UNKNOWN
I cannot determine the specific residue numbers reported for Pigeonpox virus PDE from the available literature.

**open-book** · **correct** · 4 tool calls
H72, H167
These residues were identified as key catalytic histidines in Pigeonpox LigT-like phosphodiesterase (PDE) required for the enzymatic hydrolysis of cGAMP isomers (such as 2′,3′-cGAMP and 3′,3′-cGAMP) in in vitro biochemical assays (DOI: 10.1038/s41586-024-07809-y).

**rs29 · Which amino acid residues were reported as important for the interaction between TMPRSS2 and HCoV-NL63-S2′? Context: Biolayer interferometry (BLI) affinity measurement. List the residues.**

gold Arg870, Ile867 (doi 10.1038/s41594-026-01801-y, 2026)

**closed-book** · **unknown**
UNKNOWN
I cannot determine the specific residues identified as important for the TMPRSS2 and HCoV-NL63-S2′ interaction in the biolayer interferometry study.

**open-book** · **correct** · 4 tool calls
Arg870, Ile867

These residues were identified as important determinants for the binding of the HCoV-NL63-S2′ peptide to TMPRSS2 in biolayer interferometry (BLI) affinity assays (KD ~ 500 nM). This finding comes from a study investigating TMPRSS2-mediated coronavirus spike recognition and activation (DOI: 10.1038/s41594-026-01801-y).

**rs30 · Which amino acid residues were reported as important for the interaction between MIS12C and Dsn1N? Context: ITC measurement of Dsn1(92-113) binding to MIS12C. List the residues.**

gold Trp96, Arg97, Arg98 (doi 10.1038/s41594-024-01249-y, 2024)

**closed-book** · **unknown**
UNKNOWN

**open-book** · **correct** · 2 tool calls
W96, R97, R98

These residues form the conserved WRR motif within the auto-inhibitory Dsn1(92–113) segment that mediates binding to MIS12C in ITC measurements (DOI: 10.1038/s41594-024-01249-y).

**rs31 · Which amino acid residues were reported as important for the interaction between SdnG and Substrate 4? Context: Imine trapping assay and site-directed mutagenesis. List the residues.**

gold His72, Asp24 (doi 10.1038/s41929-025-01294-w, 2025)

**closed-book** · **unknown**
UNKNOWN

**open-book** · **correct** · 2 tool calls
D24, H72

These residues form a catalytic dyad essential for activating K127 as a nucleophile to generate the covalent imine/iminium intermediate with the substrate. In imine trapping assays, H72A and D24A mutations abolished Schiff base adduct formation.

#### TYPE 3 · FINDINGS PUBLISHED IN 2026 (AFTER THE GENERATOR'S TRAINING DATA)

Correct only if both blind judges (Claude, OpenAI) label it correct against the gold finding; any other combination counts as not correct.

##### rc32 · A 2026 study examined STING and T-cells. Context: TIMER 2.0 and Kaplan-Meier Plotter analysis. What did it report?

gold High STING expression is associated with increased CD4+ and CD8+ T-cell infiltration and improved prognosis. (doi 10.1155/ogi/2210380, 2026)

**closed-book** · **unknown**
UNKNOWN

**open-book** · **unknown** · 4 tool calls
UNKNOWN

##### rc33 · A 2026 study examined ARX and PTPRD. Context: CRISPRi screen across 44 TFs. What did it report?

gold Convergent effector genes downstream of TF perturbations are enriched for neuropsychiatric disease-associated genes (SCZ/MDD). (doi 10.1038/s41586-025-09997-7, 2026)

**closed-book** · **unknown**
UNKNOWN

**open-book** · **correct** · 2 tool calls · judges: claude correct, openai correct
PTPRD is a convergent downstream effector gene regulated by ARX (and other transcription factors) that is enriched for neuropsychiatric disorder (schizophrenia/SCZ and major depressive disorder/MDD) risk genes ($P < 0.05$).

This comes from a 2026 single-cell CRISPRi (Perturb-seq) screen evaluating 44 transcription factors in human radial glia during cortical neurogenesis (DOI: 10.1038/s41586-025-09997-7). Convergent downstream effectors like PTPRD showed significant enrichment for neuropsychiatric disease-associated gene sets by Fisher's exact test.

##### rc34 · A 2026 study examined TMPRSS2 and H1H7. Context: In vitro neutralization assay using Vero E6-TMPRSS2 cells and VSV pseudotyped SARS-CoV-2. What did it report?

gold The H1H7 antibody potently inhibits SARS-CoV-2 variants by sterically blocking TMPRSS2, with an IC50 of ~65 ng/ml. (doi 10.1038/s41594-026-01801-y, 2026)

**closed-book** · **unknown**
UNKNOWN

**open-book** · **correct** · 2 tool calls · judges: claude None, openai correct
IC50 of ~65 ng/ml

This finding is reported in a 2026 study investigating TMPRSS2-mediated coronavirus spike activation (DOI: 10.1038/s41594-026-01801-y). The monoclonal antibody H1H7 potently inhibits SARS-CoV-2 infection by sterically blocking TMPRSS2 access to the early fusion intermediate conformation (E-FIC).

**rc35 · A 2026 study examined HER2 and T-DXd. Context: Clinical trial DESTINY-Breast04. What did it report?**

gold T-DXd significantly improved progression-free survival in patients with HER2-low metastatic breast cancer compared to chemotherapy (HR=0.51, P<0.001). `(doi 10.1158/1078-0432.ccr-25-1982, 2026)`

**closed-book** · **unknown**
UNKNOWN

**open-book** · **correct** · 2 tool calls · judges: claude correct, openai correct
T-DXd significantly improved progression-free survival in patients with HER2-low metastatic breast cancer compared to chemotherapy (HR = 0.51, P < 0.001).

This result is reported in a 2026 review on targeting HER2 across cancer subtypes (DOI: 10.1158/1078-0432.CCR-25-1982), which synthesizes clinical trial findings from the DESTINY-Breast04 study.

**rc36 · A 2026 study examined Rad21 and Pou5f3. Context: Hi-C analysis at 5.3 hpf zebrafish embryos. What did it report?**

gold Fountains are distinct chromatin structures enriched at enhancers that emerge following zygotic genome activation (ZGA). `(doi 10.1038/s41467-026-69105-9, 2026)`

**closed-book** · **unknown**
UNKNOWN

**open-book** · **unknown** · 6 tool calls
UNKNOWN

**rc37 · A 2026 study examined ATG5-ATG12 and ALIX. Context: LLOMe damage in ATG5 KO vs ATG8 KO HeLa cells. What did it report?**

gold The ATG5-ATG12 conjugate is essential for ESCRT recruitment; ATG8 lipidation is required for optimal, stabilized ESCRT architecture but is not strictly necessary for initial recruitment. `(doi 10.1038/s44318-025-00672-1, 2026)`

**closed-book** · **unknown**
UNKNOWN

**open-book** · **correct** · 2 tool calls · judges: claude correct, openai correct
The ATG5-ATG12 conjugate is essential for ESCRT recruitment (including ALIX), whereas ATG8 lipidation is required for optimal, stabilized ESCRT architecture but is not strictly necessary for initial recruitment.

This finding comes from a 2026 study investigating ATG8 E3-like ligase complexes in LLOMe-damaged HeLa cells (DOI: 10.1038/s44318-025-00672-1). Comparing ATG5 KO and ATG8 KO cells demonstrated that ATG5-ATG12 acts upstream to initiate ESCRT machinery recruitment to damaged lysosomes.

**rc38 · A 2026 study examined NSD2 and H3K36me2. Context: CRISPR-Cas9 knockout in NPPO-1NE and MSKPCa10 organoids. What did it report?**

gold NSD2 is required for maintaining the neuroendocrine phenotype in CRPC-NE organoids. `(doi 10.1038/s41586-025-09727-z, 2026)`

**closed-book** · **unknown**
UNKNOWN

**open-book** · **correct** · 4 tool calls · judges: claude correct, openai correct
NSD2 is required for maintaining the neuroendocrine phenotype in castration-resistant neuroendocrine prostate cancer (CRPC-NE) organoids, and its knockout drives reversion to an adenocarcinoma phenotype.

This finding comes from a study showing that CRISPR-Cas9 knockout of NSD2 in NPPO-1NE and MSKPCa10 organoids depletes H3K36me2, reversing lineage plasticity and restoring the canonical androgen receptor program (DOI: 10.1038/s41586-025-09727-z).

**rc39 · A 2026 study examined Aldotetrose and Glycolaldehyde. Context: Microkinetic simulation. What did it report?**

gold The Breslow autocatalytic cycle is kinetically suppressed at high glycolaldehyde concentrations and operates predominantly at low concentrations. (`doi 10.1021/jacsau.5c01359, 2026`)

**closed-book** · **unknown**
UNKNOWN

**open-book** · **correct** · 2 tool calls · judges: claude correct, openai correct
The Breslow autocatalytic cycle is kinetically suppressed at high glycolaldehyde concentrations and operates predominantly at low concentrations.

This finding was reported in a 2026 computational study investigating formose reaction mechanisms using molecular dynamics and DFT-based microkinetic simulations (DOI: 10.1021/jacsau.5c01359).

**rc40 · A 2026 study examined ZCWPW1-ZCWPW2 complex and LDHA. Context: L-LDH activity measured by WST-8 assay in KO vs WT testis tissue. What did it report?**

gold The ZCWPW1-ZCWPW2 complex binds to and stabilizes the enzymatic activity of LDHA, increasing lactate production. (`doi 10.1093/nar/gkag049, 2026`)

**closed-book** · **unknown**
UNKNOWN

**open-book** · **correct** · 4 tool calls · judges: claude correct, openai correct
The ZCWPW1-ZCWPW2 complex binds to and stabilizes the enzymatic activity of LDHA, increasing lactate production (with reduced L-LDH activity in knockout testis tissue).

This finding comes from a 2026 study on meiotic recombination regulation by ZCWPW2 and ZCWPW1 (DOI: 10.1093/nar/gkag049). WST-8 enzymatic assays demonstrated that loss of the complex reduces LDHA catalytic activity without altering LDHA total protein abundance.

**rc41 · A 2026 study examined NTSR1 and G protein. Context: BRET2-based dissociation/recruitment in CHO-hNTSR1 cells. What did it report?**

gold PP-001 acts as a biased allosteric agonist of NTSR1, favoring G protein (Gαq, Gαi1, GαoA, Gα13) activation over β-arrestin recruitment. (`doi 10.1016/j.apsb.2025.12.043, 2026`)

**closed-book** · **unknown**
UNKNOWN

**open-book** · **correct** · 3 tool calls · judges: claude correct, openai correct
41-68% max activation for G proteins (0% beta-arrestin recruitment)
In a 2026 study investigating the NTSR1-derived pepducin PP-001 (DOI: 10.1016/j.apsb.2025.12.043), BRET2 assays in CHO-hNTSR1 cells showed that PP-001 acts as a biased allosteric agonist that selectively promotes G protein activation (Gαq, Gαi1, GαoA, Gα13) over β-arrestin recruitment.

**rc42 · A 2026 study examined BrNiR and AcNiR. Context: Atomic resolution XFEL structures at various pH states. What did it report?**

gold All oxidised CuNiR structures exhibit a pentacoordinated T2Cu(II) catalytic site. (doi 10.1038/s41467-026-70261-1, 2026)

**closed-book** · **unknown**
UNKNOWN
I cannot reliably identify or confirm the findings of a 2026 study on BrNiR and AcNiR.

**open-book** · **correct** · 3 tool calls · judges: claude correct, openai correct
All oxidized CuNiR structures exhibit a pentacoordinated T2Cu(II) catalytic site (at 1.00–1.15 Å resolution).

This was demonstrated using damage-free, atomic-resolution serial femtosecond rotational crystallography (SF-ROX) of copper nitrite reductases from *Bradyrhizobium* (BrNiR) and *Achromobacter cycloclastes* (AcNiR) across various pH states (DOI: 10.1038/s41467-026-70261-1). The study resolved long-standing questions about CuNiR coordination architecture by confirming a conserved pentacoordinated geometry in the resting oxidized state across species.

**rc43 · A 2026 study examined SplA and Ac-YLY-AMC. Context: Fluorescence activity assay in PBS pH 7.4. What did it report?**

gold SplA allelic variant A3 shows higher activity than wildtype, while variants A2 and A7 are inactive against SplA-specific peptide substrates. (doi 10.1002/cbic.202500816, 2026)

**closed-book** · **unknown**
UNKNOWN

**open-book** · **correct** · 2 tool calls · judges: claude correct, openai correct
SplA allelic variant A3 shows higher activity than wildtype, while variants A2 and A7 are inactive against the substrate.

This finding was reported in a 2026 study investigating the substrate specificity and stability of *Staphylococcus aureus* SplA and SplB serine protease allelic variants using fluorescence-based cleavage assays in PBS pH 7.4 (DOI: 10.1002/cbic.202500816).

**rc44 · A 2026 study examined CYP7A1 and ABCB11. Context: Condition 2 vs Condition 1 media. What did it report?**

gold Condition 2 (800 pM insulin + 11 mM glucose + 100 µM FFA) induces significant steatosis and bile acid elevation. (doi 10.1038/s42003-025-09484-9, 2026)

**closed-book** · **unknown**
UNKNOWN

**open-book** · **unknown** · 7 tool calls
UNKNOWN

**rc45 · A 2026 study examined ATF1 and SMC1B. Context: In vitro EMSA binding assay. What did it report?**

gold The risk allele rs2272804 in the SMC1B promoter significantly reduces ATF1 binding affinity. (doi 10.1038/s41586-025-09964-2, 2026)

**closed-book** · **unknown**
UNKNOWN

**open-book** · **correct** · 3 tool calls · judges: claude correct, openai correct
Kd = 173.39 nM (variant rs2272804) vs Kd = 56.62 nM (reference)

In an in vitro EMSA binding assay, the meiotic aneuploidy risk allele rs2272804 in the *SMC1B* promoter reduced ATF1 transcription factor binding affinity by more than 3-fold compared to the reference sequence (Kd = 173.39 nM vs 56.62 nM). This finding comes from a 2026 study investigating genetic variation in human meiotic recombination and aneuploidy (DOI: 10.1038/s41586-025-09964-2).

**rc46 · A 2026 study examined IFN-λ and RVFV. Context: hTSC culture treated 24h prior to infection. What did it report?**

gold Pretreatment with recombinant IFN-λ (100 ng/mL) significantly inhibits RVFV protein expression (Gn and NSs) in human trophoblast stem cells. `(doi 10.1016/j.isci.2026.115584, 2026)`

**closed-book** · **unknown**
UNKNOWN
I cannot reliably identify the specific 2026 study or its findings regarding IFN-λ pretreatment and RVFV in hTSCs.

**open-book** · **correct** · 2 tool calls · judges: claude None, openai correct
Pretreatment with recombinant IFN-λ (100 ng/mL) significantly inhibits RVFV protein expression (Gn and NSs).

This was demonstrated in a 2026 study investigating Rift Valley fever virus (RVFV) infection in primary human trophoblast stem cells (hTSCs) (DOI: 10.1016/j.isci.2026.115584). Exogenous application of type III interferon 24 hours prior to viral exposure mounted a robust antiviral response that suppressed viral replication.

**rc47 · A 2026 study examined KIT and KITbp. Context: X-ray crystallography. What did it report?**

gold KITbp:KIT crystal structure validates the design approach, showing sub-angstrom backbone RMSD. `(doi 10.1038/s41467-025-67866-3, 2026)`

**closed-book** · **unknown**
UNKNOWN

**open-book** · **unknown** · 7 tool calls
UNKNOWN
